# T2G: fast and accurate transcriptome to genome BAM liftover for short and long read RNA sequencing

**DOI:** 10.64898/2026.10.20.760397

**Authors:** Danielle L. Rudler, Shafi Mahmud, Alice Cleynen, Nikolay E. Shirokikh

## Abstract

Aligning RNA sequencing reads to the transcriptome rather than the genome recovers short and junction-spanning reads that genome alignment loses or misplaces, but it leaves the alignments in transcript coordinates, which genome browsers, variant callers and most annotation tools cannot use. Existing liftover tools translate between genome assemblies, and the few that reach transcript coordinates are tied to a single quantification model or assay type. Here we describe T2G, which converts complete BAM files from transcriptome to genome coordinates while keeping CIGAR strings, alignment tags and per-base RNA modification calls intact. Across nanopore direct RNA sequencing, ribosome footprinting and paired-end Illumina RNA sequencing, aligning to the transcriptome and lifting with T2G gave higher on-target specificity and more accurate splice junctions than direct genome alignment, removed the systematic assignment of reads to processed pseudogenes, and preserved isoform identity, revealing isoform-specific modification levels that genome alignment pools together. In a reanalysis of a published RNA-binding protein experiment, T2G reduced the number of reads attributed to pseudogenes 8.5-fold compared with unspliced genome alignment. T2G is available at https://github.com/Arnaroo/T2GD as a Python reference implementation and as t2gd, a dependency-free binary for Linux, macOS and Windows that also provides a gene-level quantifier and BAM utilities toolkit.

## Introduction

Every RNA sequencing workflow must choose between genome and transcriptome alignment. Genome alignment returns the chromosomal coordinates that nearly every downstream tool expects; transcriptome alignment returns sensitivity. Mapping the same libraries with and without a reference gene model, more than a third of junction-spanning reads cannot be aligned at all without one, only about half map to the same genomic position with and without one, and for reads crossing three or more exons agreement falls below 7% (Zhao, 2014). Reads that miss their junction do not vanish. Processed pseudogenes, unspliced genomic copies of the very transcripts being sequenced, are the usual destination, and they concentrate in the highly expressed housekeeping families that dominate RNA sequencing libraries (Pei et al., 2012; Cheetham et al., 2020). Transcriptome-first pipelines have accordingly been standard since TopHat2 (Kim et al., 2013) and are close to mandatory for ribosome profiling, where footprints of roughly 30 nucleotides cannot be placed across a junction unaided (Ingolia et al., 2009). The price is paid at the end: the output BAM is written in transcript coordinates and is unusable in a genome browser, in variant calling, or in any tool keyed to chromosomes. Direct RNA sequencing sharpens the problem, since long reads span many exons at once and carry per-base modification tags that must survive the journey intact (Garalde et al., 2018; Workman et al., 2019).

No existing tool closes the gap. Liftover between genome assemblies is mature (Kent et al., 2002; Zhao et al., 2014), and levioSAM2 (Chen et al., 2024) shows that CIGAR strings and SAM tags can be carried faithfully through a lift, but an assembly-to-assembly chain is a nearly colinear mapping between two versions of one coordinate system. A transcript-to-genome mapping is not: it must introduce intron skips absent from the source alignment, for every read, driven by an annotation rather than a chain file. Tools working natively in transcript coordinates, including GenomicRanges (Lawrence et al., 2013), AGouTI (Kosiński and Żywicki, 2023) and our own R2Dtool (Sethi et al., 2024), move intervals rather than alignments. Two tools perform the conversion itself, both within narrow bounds: rsem-tbam2gbam (Li and Dewey, 2011) is tied to RSEM’s probabilistic quantification model and was validated on short paired-end Illumina data, and the mergeBAM module of BASALkit (Xu et al., 2025) is confined to short-read base-conversion data and makes no provision for tag fidelity.

Here we present T2G, a general, annotation-driven converter that treats the transcriptome-aligned BAM as the primary object, preserves CIGAR structure together with the full tag payload including base-modification tags, and serves short and long reads alike (Fig. 1).

**Figure 1.**
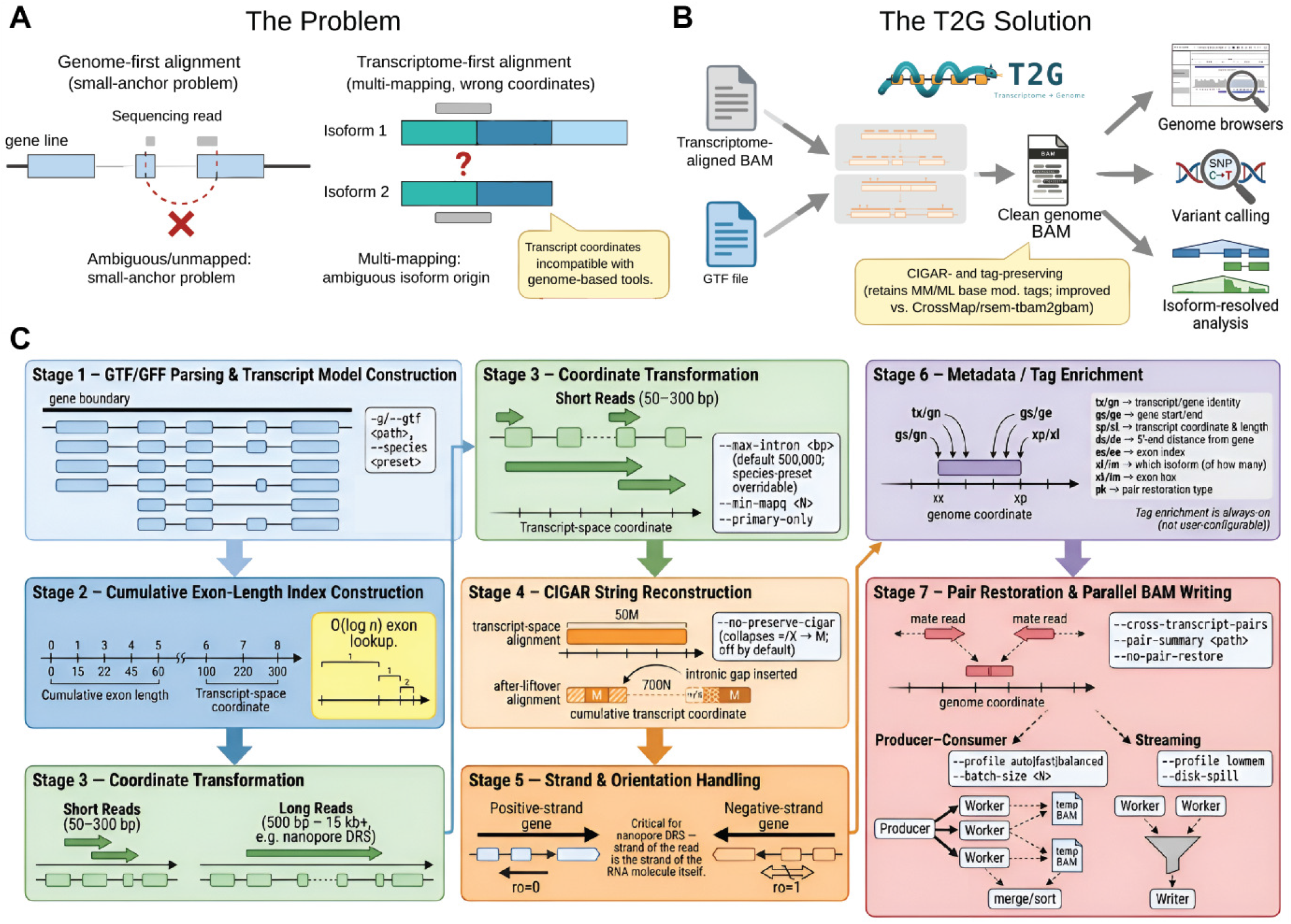
T2G converts transcriptome-aligned BAM files to genome coordinates. (**A**) Genome-first alignment suffers the small-anchor problem, in which junction-spanning reads may be dropped, truncated or misplaced; transcriptome-first alignment multi-maps across isoforms and returns transcript coordinates incompatible with downstream tools. (**B**) Transcript coordinates are mapped through the annotated exon structure, intron skips are inserted as N operations, and CIGAR strings, alignment tags and base-modification tags are carried across unchanged. (**C**) Conversion pipeline, from annotation parsing and exon-length indexing through coordinate transformation, CIGAR reconstruction, strand handling and tag enrichment to pair restoration and parallel BAM writing.

## Methods

### T2G algorithm and implementation

T2G parses GTF/GFF annotations into transcript models that record exon boundaries, strand information and, crucially, the gene boundaries across all isoforms, then maps each alignment to genomic coordinates by binary search through cumulative exon lengths (*O*(log *n*); Fig. 1C). Genomic CIGAR strings are reconstructed preserving the M (match/mismatch), = (sequence match), and X (sequence mismatch) distinction and inserting validated intron N operations, including for long reads with many intron spans. Negative-strand alignments are reverse-complemented with base qualities reversed. Because junction-spanning reads are contiguous matches in transcript space, junction placement is resolved deterministically from the annotation at the lift rather than inferred from a few anchoring nucleotides, which avoids the small-anchor problem that makes genome alignment of ultra-short and short junction-spanning reads prone to failure. The same pipeline serves ultra-short (15 to 40 nt, e.g. footprinting), short (50 to 300 bp) and long (500 bp to 15 kb and above) reads. By default T2G warns for introns larger than 100 kb and rejects reads implying introns larger than 500 kb; species presets adjust the rejection threshold (for example yeast 5 kb, plant 750 kb) and all thresholds are configurable.

All input tags, including base modifications (MM/ML), are preserved, and T2G adds the positional and mapping tags in Table 1, all lowercase two-character tags, the space the SAM specification reserves for end-user applications. In paired-end mode, pairs and mate tags are restored during liftover, including for multi-mapping pairs. Two parallelisation strategies, producer–consumer (workers convert batches and write temporary BAMs that are merged) and streaming (a single sequential writer), support high-throughput and memory-constrained use respectively, and a parsed-GTF cache and disk spill under a user-set memory budget are available. T2G is provided as a readable Python reference implementation and as t2gd, a compiled binary built from a separate D implementation of the same method.

**Table 1.** Tags written by T2G.

| Tag(s) | Content |
| --- | --- |
| tx, gn | Transcript and gene identifiers |
| gs, ge | Genomic bounds of the alignment (gs leftmost, 0-based; ge rightmost, half-open) |
| xp, xl | Genomic span including introns (xp); exonic length excluding introns (xl) |
| xi, im | Intron-spanning flag (xi); largest intron crossed (im, written only when xi = 1) |
| ds, de, es, ee | Gene-relative positions from the 5' and 3' ends, in the direction of transcription |
| ro | Orientation (1 forward, -1 reverse) |
| pk | Pair-restoration outcome (0 singleton, 1 pair on one transcript, 2 pair across transcripts) |
| NH | Alignment multiplicity |

### Auxiliary toolkit

t2gd ships as a single static binary of 36 subcommands (Table 2), with an optional GTK graphical interface exposing the same functionality flag for flag. count is a featureCounts-equivalent gene-level quantifier operating directly on a genome-coordinate BAM and GTF, and is used for all gene-detection comparisons below; inject-summary attaches Oxford Nanopore run metadata to existing BAMs; and tagcheck audits SEQ and base-modification tag coverage by alignment class.

**Table 2.** t2gd subcommand groups.

| Group | Subcommands |
| --- | --- |
| Convert & inject | convert, inject-summary |
| Inspect | view, flagstat, idxstats, stats, depth, coverage, bedcov, head, flags, samples, quickcheck, validate, checksum, tagcheck |
| Extract | fastq, fasta, faidx, dict |
| Manipulate | sort, index, calmd, fixmate, collate, merge, cat, encode |
| Surgery | addreplacerg, filter, subsample, split, reheader, dedup |
| Counting | count |

### References and datasets

Human analyses used the GRCh38 primary assembly and GRCh38.115 annotation (Homo sapiens.GRCh38.dna.primary assembly, Ensembl release 115), with a complementary transcriptome reference extracted by gffread (Pertea and Pertea, 2020). The scalability benchmark used Ensembl release 115 references for rat (Rattus_norvegicus.GRCr8.cdna.all) and yeast (Saccharomyces_cerevisiae.R64-1-1.cdna.all) and Ensembl Plants release 62 for *A. thaliana* (TAIR10/Araport11) (Table 3). An additional *S. cerevisiae* S288C reference genome (R64-5-1, release 20240529) was downloaded from the Saccharomyces Genome Database (SGD; http://sgd-archive.yeastgenome.org); this is the ORF-based reference, and it is used only for the auxiliary toolkit comparison.

**Table 3.**
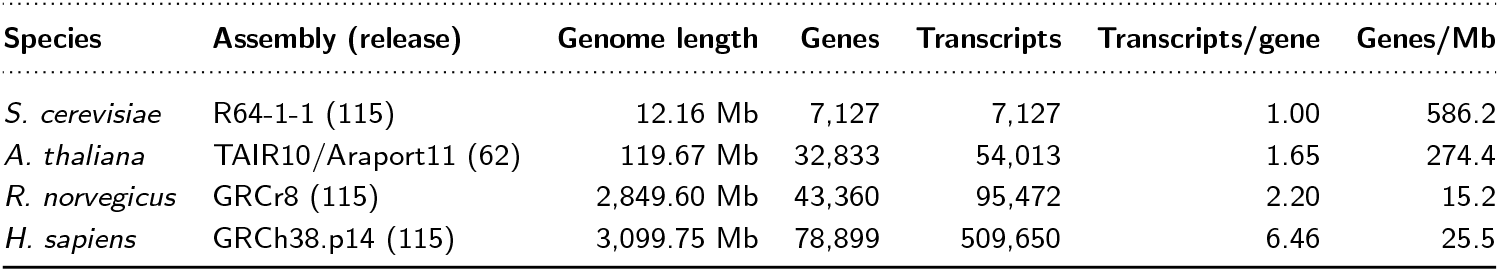
Genome and annotation statistics for the runtime benchmark species (gene and transcript feature lines per GTF; genome length from the NCBI assembly report, primary assembly plus organelles).

Direct RNA sequencing (DRS) data were generated for this study (BioProject PRJNA1512710). HEK293T cells were grown in DMEM with 10% fetal bovine serum and 1*×* antibiotic-antimycotic at 37 ^*°*^C and 5% CO_2_, scraped, lysed through an 18–31G needle and clarified at 12,000 *g* for 5 min at 4 ^*°*^C. Total RNA was extracted with the PureLink RNA Mini Kit (Invitrogen) and quantified with the Qubit RNA High Sensitivity assay. 2 *μ*g total RNA was prepared with the Direct RNA Sequencing Kit (SQK-RNA004, ONT) using Induro reverse transcriptase and sequenced on a MinION. All other datasets are public: 2*×*100 bp HEK293T mRNA-seq (SRR31348821 and replicate SRR31348820; Damodaran et al., 2025), HEK293 ribosome footprints (SRR493747; Ingolia et al., 2012), paired-end libraries from rat (SRR33313115), yeast (SRR18592149) and *A. thaliana* (ERR16142461) for scalability, and a 2*×*150 bp POND-seq library (SRR37350470; Hu et al., 2026) for the RNA-binding protein reanalysis.

### Alignment pipelines

DRS reads were basecalled with Dorado 1.3.0 (rna004_130bps_sup@v5.2.0, with m6A, pseudouridine, m5C and 2^*′*^-O-methylation calling and poly(A) estimation). Two alignment passes were run from the same basecalled reads: a genome-direct alignment that used Dorado’s built-in minimap2 (-x splice -k14 -N10) against the GRCh38 primary assembly, and a T2G-liftover that used minimap2 (-x map-ont -k14 -N10) against the extracted transcriptome followed by t2gd convert (v1.17.52, --validate --no-pair-restore). Paired-end reads were trimmed with Trim Galore v2.2.0 and aligned with bowtie2 2.5.5 (Langmead and Salzberg, 2012) to the genome and, separately, to the transcriptome followed by T2G with pair restoration (v1.17.52, --pair-streaming, pair restoration on). Because bowtie2 is not splice-aware, using it in both arms isolates reference choice from aligner choice; STAR 2.7.11b (Dobin et al., 2013; GTF-guided index, --sjdbOverhang 100 --quantMode TranscriptomeSAM --outSAMunmapped Within) was added as a splice-aware genome baseline. Footprint reads were linker-clipped and first-base-trimmed with cutadapt v5.2 (Martin, 2011), retaining 46,976,676 of 52,070,666 reads, and aligned with bowtie2 2.5.4 to genome and transcriptome; the STAR arm was configured so that splice-awareness was the only difference from bowtie2 (--alignEndsType EndToEnd, --outFilterMultimapNmax 1000, --seedSearchStartLmax 20, --outSAMunmapped Within). POND-seq pairs were processed through the same three pipelines. Every arm retained unmapped reads so that all arms of a comparison share one denominator. The STAR footprint arm and the paired-end STAR mapping rates are from realignments of the same reads to the same Ensembl 115 index. DRS, paired-end and footprint conversions used t2gd v1.17.52; the runtime benchmark and the POND-seq reanalysis used v1.19.0, whose conversion record stream is byte-identical to the released v1.19.1, which was used for the samtools comparison.

### Benchmark analyses

#### On-target classification

For DRS and paired-end reads, each read’s CIGAR was walked, consecutive M/D operations were merged into exonic blocks (breaking only at N, since nanopore reads carry frequent short indels that must not be mistaken for splicing), and a read was classified as on-target, rRNA or off-target by majority (*≥*50%) overlap of its aligned span with exon and rRNA-biotype (rRNA, Mt_rRNA, rRNA_pseudogene) interval sets built from the GRCh38.115 GTF. rRNA is reported as its own category, since incomplete rRNA depletion is expected contamination rather than mRNA signal. For footprints, which are short single-block alignments, the alignment start position was tested against merged exon intervals. This simpler rule was checked rather than assumed: across all 46,210,481 mapped STAR footprint alignments there was no S operation, so the reported start is the true read start, and only 468,039 (1.01%) carried an N operation. Running both rules over the same BAMs shifted on-target by about 0.1 percentage point (STAR 7.04% by start position versus 7.15% by block-walk; bowtie2 11.91% versus 12.03%), far below the between-method differences reported. Start-position figures are used throughout for footprints. **Read fate and splice junctions**. Reads were classified by their status in each pair of pipelines (mapped in both, unmapped in both, or mapped in only one). For paired-end T2G-lifted mates spanning at least one annotated intron, the bowtie2 genome-direct fate was further classified as unmapped, same locus (mapped nearby but unable to bridge the intron) or different locus. For DRS, every distinct splice junction called by each pipeline was evaluated for canonical splice-site dinucleotides, match to an annotated transcript junction, and support by at least two reads.

#### Pseudogene placement

A genome-wide scan identified non-spliced, on-target T2G-lifted mates whose bowtie2 placement clustered elsewhere (2 kb bins). At the resulting *EEF1A1* locus, depth was computed with bcftools mpileup/call (Danecek et al., 2021) on region-restricted extracts from all three pipelines, as mean depth over the pseudogene *EEF1A1P5* (a single, fully exonic 1,388 bp block) and over the true *EEF1A1* gene, both across its full 36,279 bp genomic span and restricted to its 9,402 bp of annotated exon.

#### Isoform-resolved modification calls

At chr1:1,375,156 (*AURKAIP1*), per-read m5C calls were taken from the Dorado MM/ML tags carried into each pipeline’s BAM, and a base was counted as modified at a call probability of at least 0.5. In the T2G-lifted BAM, reads were grouped by their assigned transcript (tx tag); in the genome-direct BAM, all covering reads were pooled.

#### Footprint multi-mapping

To test whether T2G’s footprint specificity simply reflects discarding multi-mapped reads, transcriptome-side reads were realigned with bowtie2 -k2 to recover each multi-mapped read’s tied alternative alignment (rather than relying on the XS score-tie proxy available from the original -k1 run). Both alignments were translated to genome coordinates via a transcript-to-genome map built from GTF exon records, and genomic overlap was tested. The translation logic was validated against t2gd’s own output on the (transcript, position) pairs t2gd used: 12,395,644 of 12,395,644 exact matches.

#### Gene-level detection

t2gd count (exon-level, gene_id grouping, unstranded) was run on all three genome-coordinate BAMs of each comparison, with a gene detected at more than 5 assigned reads and detected genes split by GTF gene_biotype into real (protein-coding plus lncRNA), pseudogene (any pseudogene biotype) and other. Multi-mapping reads were counted once at their primary alignment (--multi-map all) rather than discarded, because bowtie2 run with -k1 reports one alignment per read without a multi-mapping flag; under the quantifier’s default, the bowtie2 arm would silently count its multi-mappers while STAR would have 43.2 million of its footprint multi-mappers discarded. As a control, the STAR arm was also counted with multi-mapped reads discarded. POND-seq fragments were counted per pair (--count-read-pairs), since the unit of evidence for an RNA-binding protein binding a transcript is a fragment; all 509,650 @SQ names resolved against the GTF. The pseudogene definition was varied (the published biotype set, a broader and a narrower set) as a sensitivity check.

#### Runtime scalability

Runtime was measured for t2gd v1.19.0 convert on nested subsets of 100,000 to 1,000,000 primary-mapped read pairs per species, at 8 threads, 5 replicates, with the parsed-GTF cache disabled (cold) and pre-populated (warm), sequentially on an exclusively allocated node of the Kaya HPC cluster (2*×* AMD EPYC 9474F, 1.5 TB RAM), and fitted as wall = *a* + *b ·* pairs.

#### Toolkit benchmarks

Four samtools-parity operations (view -c, flagstat, depth and sort) were timed for t2gd v1.19.1 against samtools 1.21 (Danecek et al., 2021) on nine BAM files (Fig. 7): the genome-direct DRS BAM (drs hek); paired-end libraries from human (hek pe), rat (rat pe) and *A. thaliana* (ath pe); the yeast library aligned to the SGD ORF and Ensembl cDNA references (yeast orf, yeast cdna); and three synthetic BAMs of increasing size, together spanning a 66-fold range in view -c work. Each cell was timed in 3 replicates on a quiescence-gated AMD EPYC 9474F node, with samtools re-timed within the same jobs. view -c, flagstat and sort were given 8 threads in both tools (-@/-t). samtools depth has no thread option, so depth was timed with t2gd at 8 threads and with both tools at one thread; it was not run on three datasets for which no coordinate-sorted BAM was produced. t2gd count was compared with featureCounts 2.1.1 (Liao et al., 2014) on a synthetic dataset with ground-truth gene assignment and on real paired-end libraries (t2gd v1.17.52, 8 threads, 2 replicates).

#### Liftover validation and performance

DRS conversion was run in validation mode (t2gd convert --validate), which reports every failed record and its cause, and t2gd tagcheck counted SEQ and MM/ML coverage by alignment class (primary, secondary, supplementary) in the genome-direct and T2G-lifted DRS BAMs. The compiled binary and the Python reference implementation were compared record by record on QNAME, FLAG, RNAME and POS under matched settings, and timed on the same cold-cache input with tag recovery disabled. Peak memory was measured with disk spill under a 1,024 MB budget.

## Results

### T2G enables transcriptome analysis at the genome scale

#### Long reads

On 6,095,970 HEK293T direct RNA reads, genome-direct spliced alignment mapped 96.0% of reads and transcriptome alignment followed by T2G 93.9%, but T2G alignments were exclusively exonic and correspondingly more precise: 98.84% of mapped reads were on-target, against 94.65%, with off-target placement eliminated (0.00% against 4.23%; Fig. 2A). The difference narrowed among reads mapped by both pipelines (98.8% against 97.0%), so it is driven mainly by reads only genome-direct places. Those reads (2.4%) outnumber T2G-only reads (0.3%) roughly 7:1, reflecting T2G’s restriction to annotated transcripts. The same pattern holds at junction level (Fig. 2B): T2G junctions were 99.9% canonical and 82.4% carried two or more reads, against 72.0% canonical and 32.8% clearing that threshold for genome-direct alignment, which called nearly three times as many distinct junctions (628,609 against 225,044).

**Figure 2.**
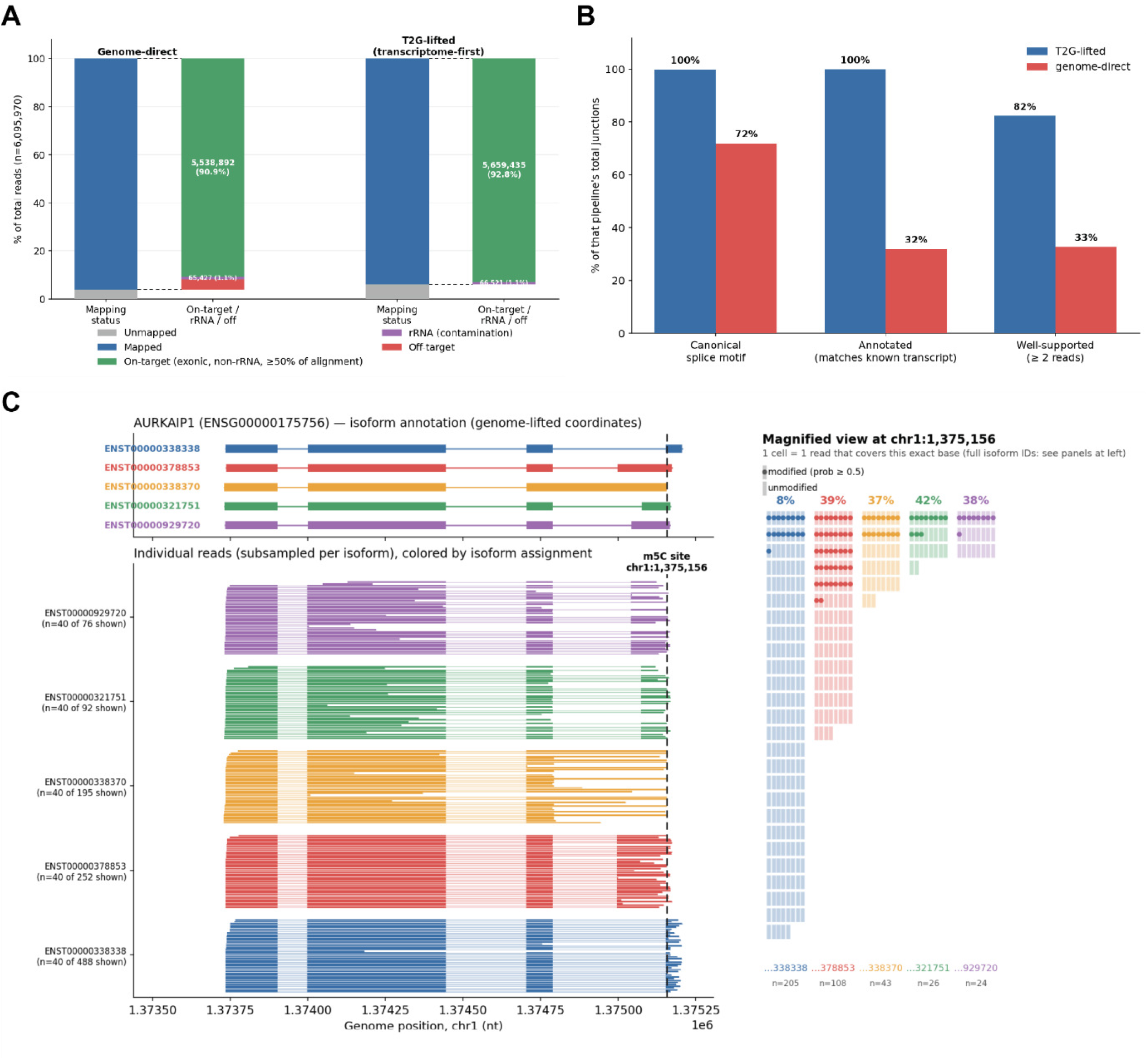
Direct RNA sequencing. (**A**) Mapping status and on-target/rRNA/off-target composition, genome-direct versus T2G-lifted, as a percentage of all 6,095,970 reads. (**B**) Percentage of each pipeline’s distinct junctions that are canonical, annotated and supported by at least two reads. (**C**) Isoform-resolved m5C stoichiometry at *AURKAIP1* chr1:1,375,156: four of five isoforms agree (37 to 42%) while ENST00000338338 does not (8.3%).

Isoform identity also survives the lift, which genome-direct alignment cannot offer. Nanopore modification callers operate in transcript space (Pratanwanich et al., 2021; Acera Mateos et al., 2024), where modification stoichiometry has been shown to differ between isoforms (Lorenz et al., 2020; Gleeson et al., 2025). At an m5C site in *AURKAIP1* (chr1:1,375,156) covered by five annotated isoforms, four agreed on stoichiometry (37 to 42% modified), while ENST00000338338, the most abundant at this site, was modified in only 8.3% of its 205 reads (Fig. 2C). Genome-direct alignment reports one pooled value (23.4%, 95 of 406 reads), and the difference disappears.

#### Paired-end short reads

On 224,114,274 HEK293T mates, the transcriptome-first route placed 96.6% of mates, against 90.2% for STAR and 82.4% for bowtie2 aligned directly to the genome. That placement rate is bowtie2’s against the transcriptome, since T2G aligns nothing. What T2G contributes is the lift, and it is close to free: 216,456,716 of 216,456,971 records survive it, a loss of 255 records (0.000118%). What the lift buys is position where T2G-lifted had the highest on-target specificity of the three (99.95%, against 97.4% and 96.0%). Essentially all (99.87%) of T2G’s sensitivity advantage over bowtie2 is attributable to junction-spanning mates, and for such mates bowtie2 more often produced a confidently wrong placement elsewhere (34.1%) than a nearby partial one (25.1%).

The mechanism is visible at *EEF1A1* (Fig. 3A). bowtie2 has no intron-skip (N) operation, so for a pair genuinely spanning an *EEF1A1* junction the true locus cannot be represented at all, leaving the processed, intronless pseudogene *EEF1A1P5* as the only site admitting a complete alignment. bowtie2 accordingly reports 78,320*×* mean exonic depth at the non-expressed pseudogene against 5,274*×* at the real gene, an inversion any naive genome-direct variant-calling or expression pipeline would read backwards, while STAR (13,997*×* against 17,069*×*) and T2G (2,384*×* against 23,478*×*) both favour the expressed locus (Fig. 3B). Transcript space represents the true transcript and the pseudogene as equally simple contiguous sequences, so the two compete on unbiased footing before the correct spliced placement is lifted back.

**Figure 3.**
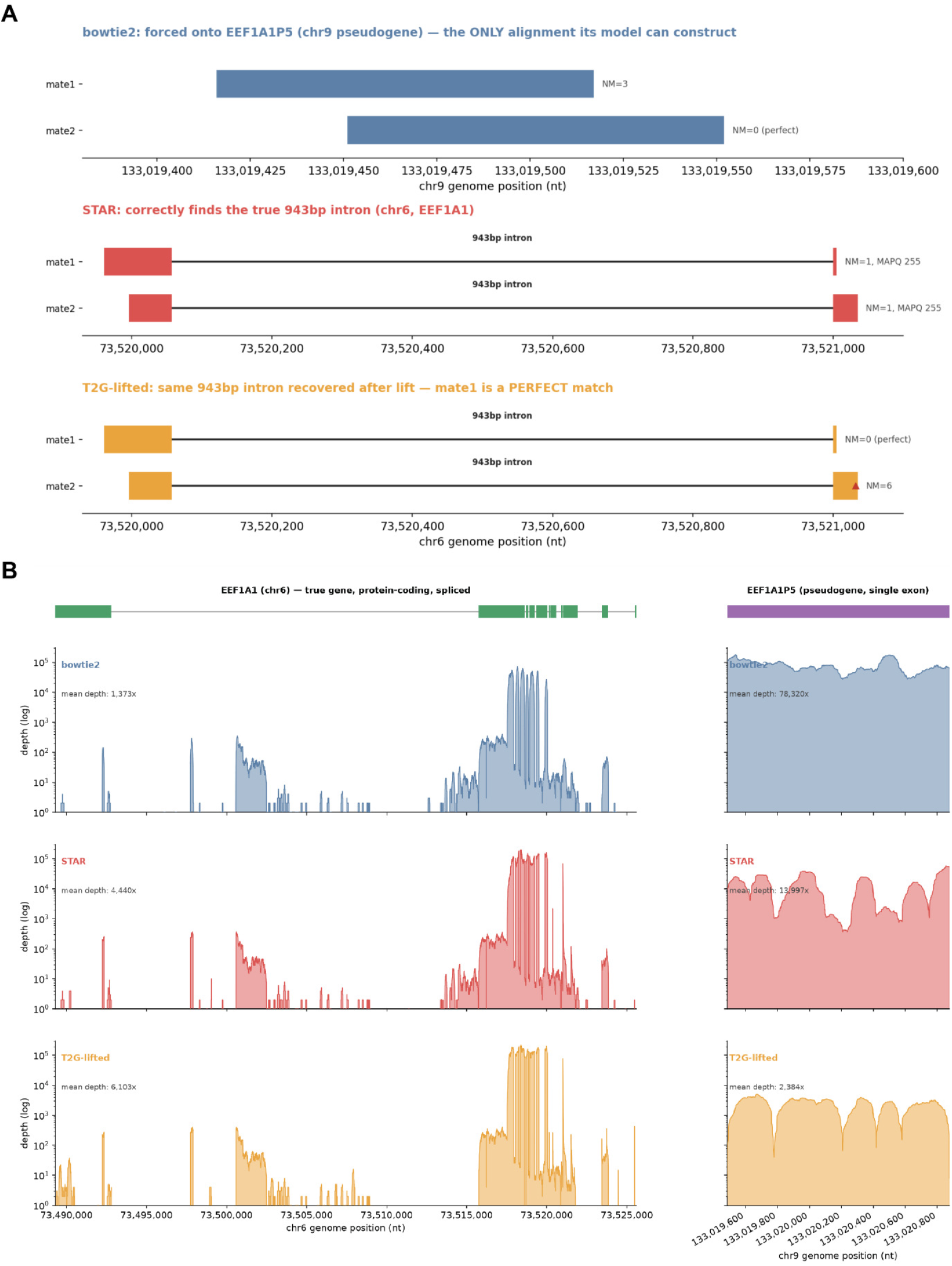
Paired-end RNA-seq. (**A**) One read pair spanning an *EEF1A1* junction: bowtie2 can only align it to the processed pseudogene *EEF1A1P5*, whereas STAR and T2G-lifted recover the spliced placement. (**B**) Read depth (log scale) across *EEF1A1* (chr6) and *EEF1A1P5* (chr9) for each pipeline.

#### Ribosome footprints

On 46,976,676 ultra-short footprint reads, both genome-direct methods mapped far more reads than transcriptome-first alignment (96.7% for bowtie2 and 98.4% for STAR, against 26.4%, T2G being limited to reads with a transcriptome hit), but that larger pool was overwhelmingly off-target: on-target exonic non-rRNA reads made up 11.9% of bowtie2 output and only 7.0% of STAR’s, against 90.5% for T2G (Fig. 4A). Genuinely spliced alignments are rare at this read length (1.0% of STAR’s mapped reads), so STAR’s extra reads are diluted by off-target and rRNA placements. In absolute terms T2G still recovered the most on-target footprints (11,217,046, against 5,408,503 for bowtie2 and 3,255,016 for STAR). This is not achieved by discarding multi-mappers: 76.8% of transcript-space multi-mapped footprints reflect genuine multi-locus genomic ambiguity that persists under any alignment strategy.

**Figure 4.**
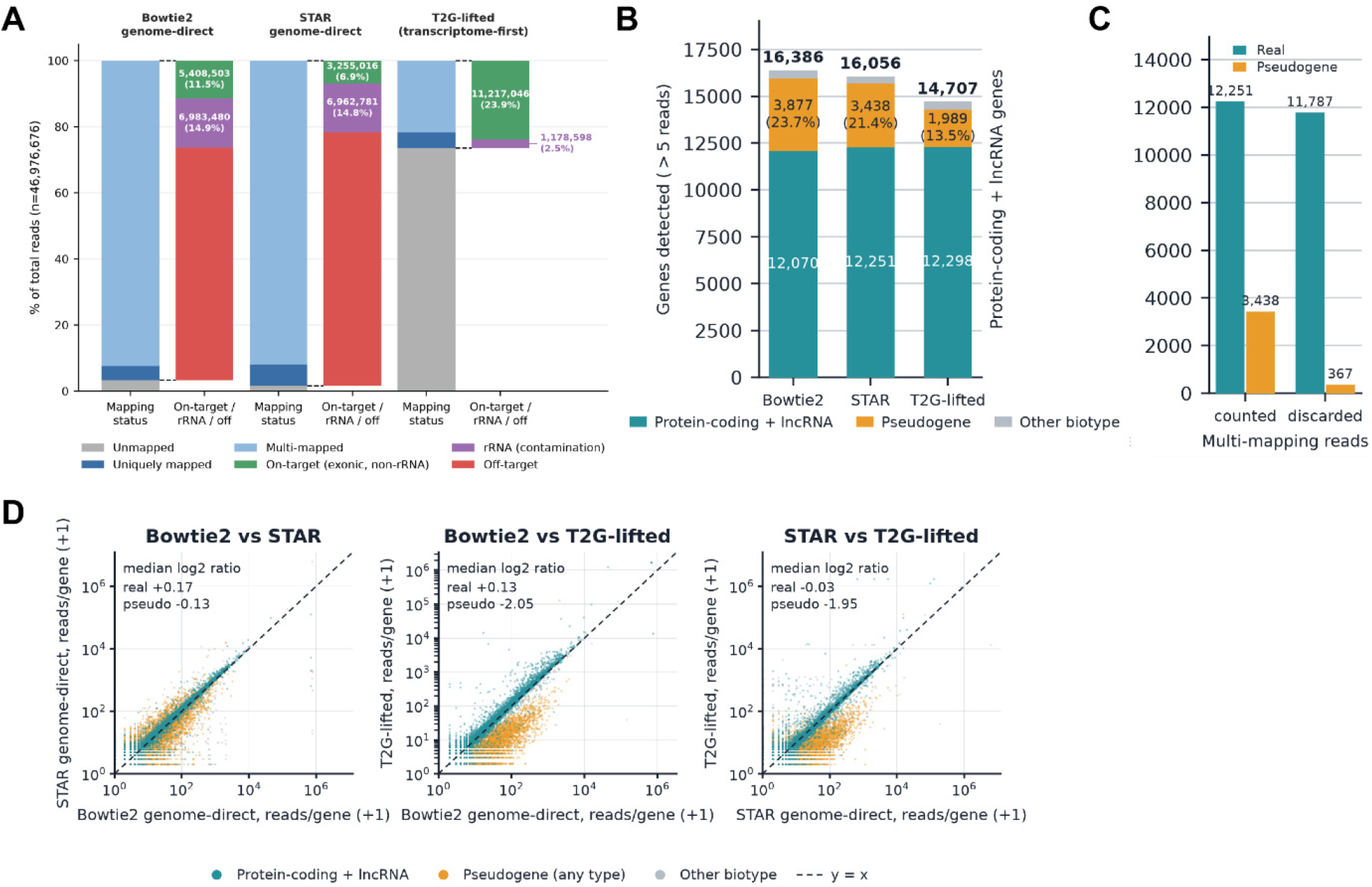
Ribosome footprinting. (**A**) Mapping status and on-target/rRNA/off-target composition, as a percentage of all 46,976,676 reads. (**B**) Genes detected (*>* 5 assigned reads) by biotype. (**C**) STAR arm counted with multi-mapped reads counted or discarded. (**D**) Per-gene read counts pairwise between pipelines, coloured by biotype; insets give median log_2_ ratios over genes with more than 5 reads in both.

Gene-level detection was not sacrificed: 12,298 protein-coding and lncRNA genes for T2G against 12,070 and 12,251, from roughly 3.7-fold fewer mapped reads, with markedly lower pseudogene contamination (13.5% of detected genes, against 23.7% and 21.4%; Fig. 4B). The near-identical contamination of two structurally different genome aligners identifies this as a reference-choice problem. Discarding multi-mapped reads in the STAR arm collapsed its pseudogene tally from 3,438 to 367 while its real-gene count barely moved (12,251 to 11,787; Fig. 4C), and per-gene counts show T2G assigning pseudogenes roughly four-fold fewer reads than either genome aligner (median log_2_ ratio *−*2.05 and *−*1.95) while real genes are essentially unmoved (Fig. 4D).

### Reanalysis of a published RNA-binding protein experiment

To test whether placement differences change biological calls, we reanalysed a published POND-seq experiment (Hu et al., 2026), in which a read assigned to a pseudogene rather than its parent gene is a wrong binding call. At more than 5 assigned reads, unspliced genome alignment attributed 2,408 detected genes to pseudogenes against 282 for T2G-lifted, a factor of 8.5, while splice-aware STAR recovered most but not all of that gap at 415 (Table 4; Fig. 5A). bowtie2 mapped 1.61 times as many pairs as the T2G-lifted arm yet assigned 27% fewer reads to genes, the signature of dispersal into paralogues rather than greater sensitivity. The effect is not a threshold artefact: across a 200-fold threshold range the T2G-lifted and STAR pseudogene shares wash out, whereas bowtie2’s barely moves (Table 4; Fig. 5B). The excess is almost entirely processed pseudogenes (2,184 against 208), the intronless retrocopies to which an unspliced aligner sends junction-spanning reads, while intron-bearing unprocessed pseudogenes show almost none (66 against 17). The result was robust to the pseudogene definition (8.5*×* to 10.5*×*), and the STAR index’s undersized junction overhang for 150 bp reads, if anything, narrows the true T2G–STAR gap.

**Table 4.**
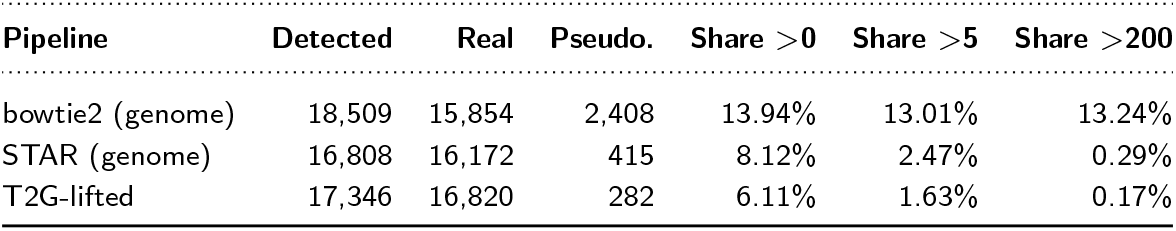
Gene detection by biotype on POND-seq SRR37350470 (13,007,301 trimmed pairs, fragment counting). Detected, real and pseudogene counts are at *>* 5 assigned reads; the last three columns give the pseudogene share of detected genes at detection thresholds of *>* 0, *>* 5 and *>* 200 assigned reads.

| Pipeline | Detected | Real | Pseudo. | Share >0 | Share >5 | Share >200 |
| --- | --- | --- | --- | --- | --- | --- |
| bowtie2 (genome) | 18,509 | 15,854 | 2,408 | 13.94% | 13.01% | 13.24% |
| STAR (genome) | 16,808 | 16,172 | 415 | 8.12% | 2.47% | 0.29% |
| T2G-lifted | 17,346 | 16,820 | 282 | 6.11% | 1.63% | 0.17% |

**Figure 5.**
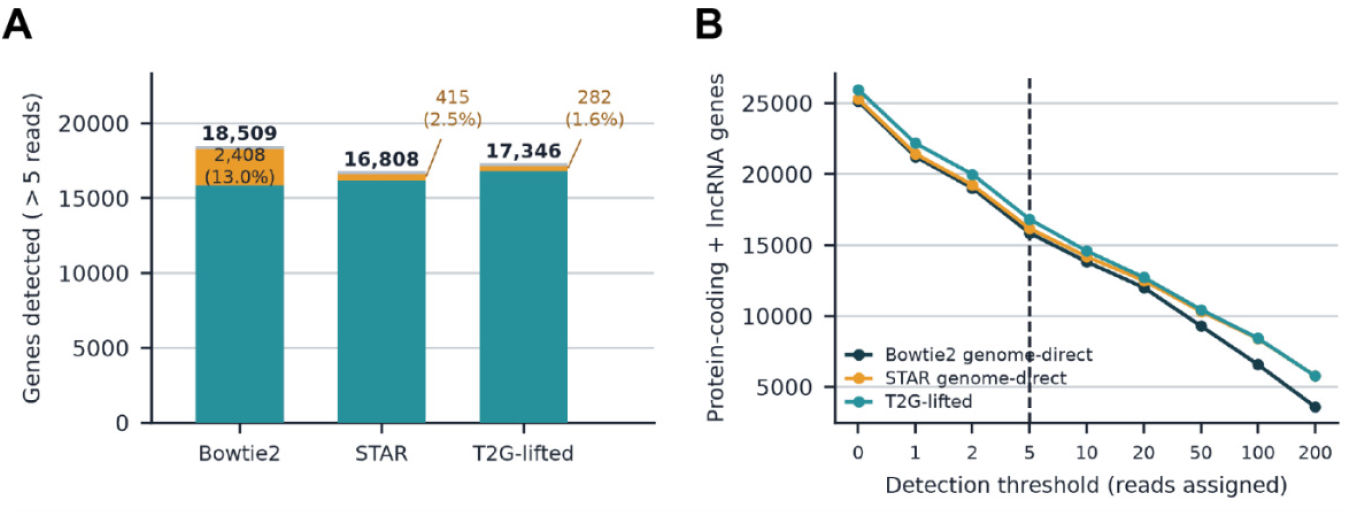
Reanalysis of POND-seq SRR37350470. (**A**) Genes detected at *>* 5 assigned reads by biotype for bowtie2 genome-direct, STAR genome-direct and T2G-lifted. (**B**) Protein-coding plus lncRNA genes detected across a threshold sweep.

### T2G is accurate, fast and efficient

#### The liftover is faithful

Run in validation mode on DRS data, T2G converted 44,525,490 of 44,525,684 mapped alignments; the 194 failures all exceeded the intron-size threshold and were reported as such. t2gd tagcheck found 100% of base-modification tags preserved across primary, secondary and supplementary alignments. Aligners such as minimap2 do not repeat modification tags on secondary and supplementary records;

T2G recovers them from the primary alignment of the same read, lifting the number of tag-bearing alignments from 444,195 to 1,935,033. On matched settings the compiled binary and the Python reference place every read identically on QNAME, FLAG, RNAME and POS; t2gd additionally retains base-modification and poly(A) tags that the reference discards.

#### Speed and memory

With tag recovery disabled so that both implementations do the same work, t2gd on eight threads converted 2.1 million records 10.2 times faster than the Python reference. With disk spill under a 1,024 MB budget, 12,000,005 records (5.02 GB) converted within 1,525 MB peak resident memory on commodity desktop hardware. Across four species spanning 12.2 Mb to 3.10 Gb of genome, runtime was affine in read count, wall = *a* + *b ·* pairs with *R*^2^ *≥* 0.9986 in every species, and the per-read term *b* spanned only 1.11-fold (54.8 to 60.8 *μ*s per pair; Fig. 6A). The between-species difference sits in the constant *a*, which is governed by annotation complexity rather than genome size: human, whose genome exceeds rat’s by 9%, carries 5.3-fold more transcripts and a 5.4-fold larger constant (24.7 s against 4.6 s; Fig. 6B–C). Caching the annotation cut human’s constant to 7.3 s (Fig. 6A, open symbols), identifying annotation ingest as the mechanism.

**Figure 6.**
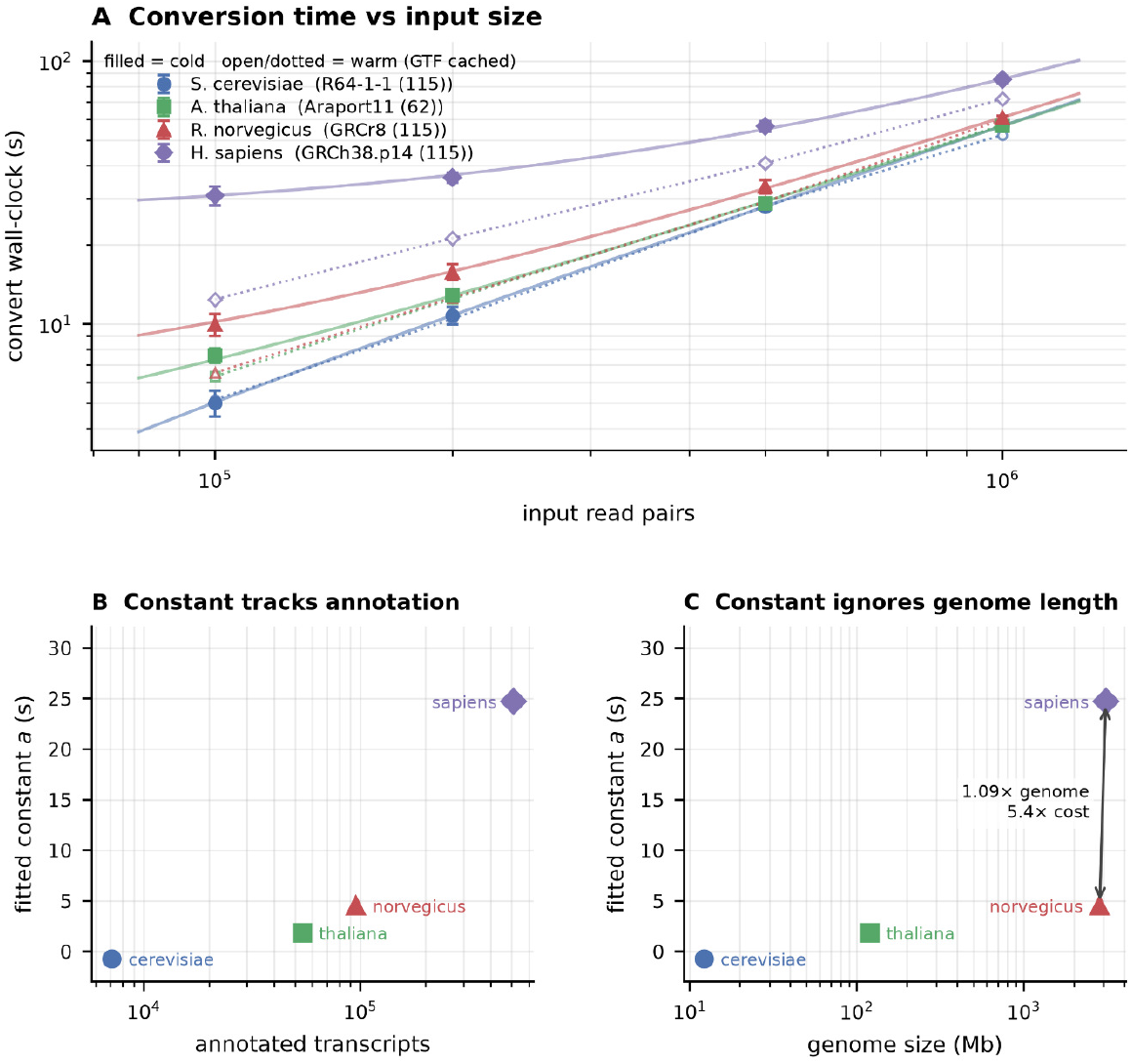
Conversion runtime (t2gd v1.19.0, 8 threads). (**A**) Wall time against input read pairs for four species (mean *±* s.d. of 5 replicates; log–log axes). Filled symbols and solid lines, cold arm (annotation parsed from disk); open symbols and dotted lines, warm arm (annotation cached). Lines are affine fits, wall = *a* + *b ·* pairs. (**B, C**) Fitted constant *a* against annotated transcript count (B) and genome size (C). The rat and human genomes differ 1.09-fold in size but their constants differ 5.4-fold, in line with their 5.3-fold difference in transcript count.

#### An accompanying toolkit

Because count is the quantifier behind every gene-detection result above, we checked it against featureCounts 2.1.1. On synthetic data with ground-truth gene assignment the two tools gave identical counts for all 37,936 genes, and on real rat and human paired-end data they agreed exactly on 80 to 84% of genes, with count consistently assigning slightly fewer reads; count was 1.4 to 4.2 times slower at 8 threads, since it does not scale with thread count. The BAM-handling subcommands were benchmarked against samtools 1.21 on nine datasets (Fig. 7). With both tools at 8 threads, t2gd was faster in every comparison: view -c by 1.26 to 2.01*×* (Fig. 7A), flagstat by 1.20 to 1.83*×* (Fig. 7B) and sort by 1.07 to 3.45*×* (Fig. 7D). The narrowest flagstat and sort margins were on the smallest BAM, where fixed start-up cost is least amortised. samtools depth is single-threaded, so most of t2gd’s 1.80 to 5.16*×* advantage at 8 threads reflects parallelism; with both tools at one thread, t2gd remained 1.27 to 2.09*×* faster (Fig. 7C). Together these subcommands cover operations otherwise reached through samtools (Bonfield et al., 2021; Danecek et al., 2021) and featureCounts, in the manner of sambamba (Tarasov et al., 2015) and elPrep (Herzeel et al., 2019), without external dependencies.

**Figure 7.**
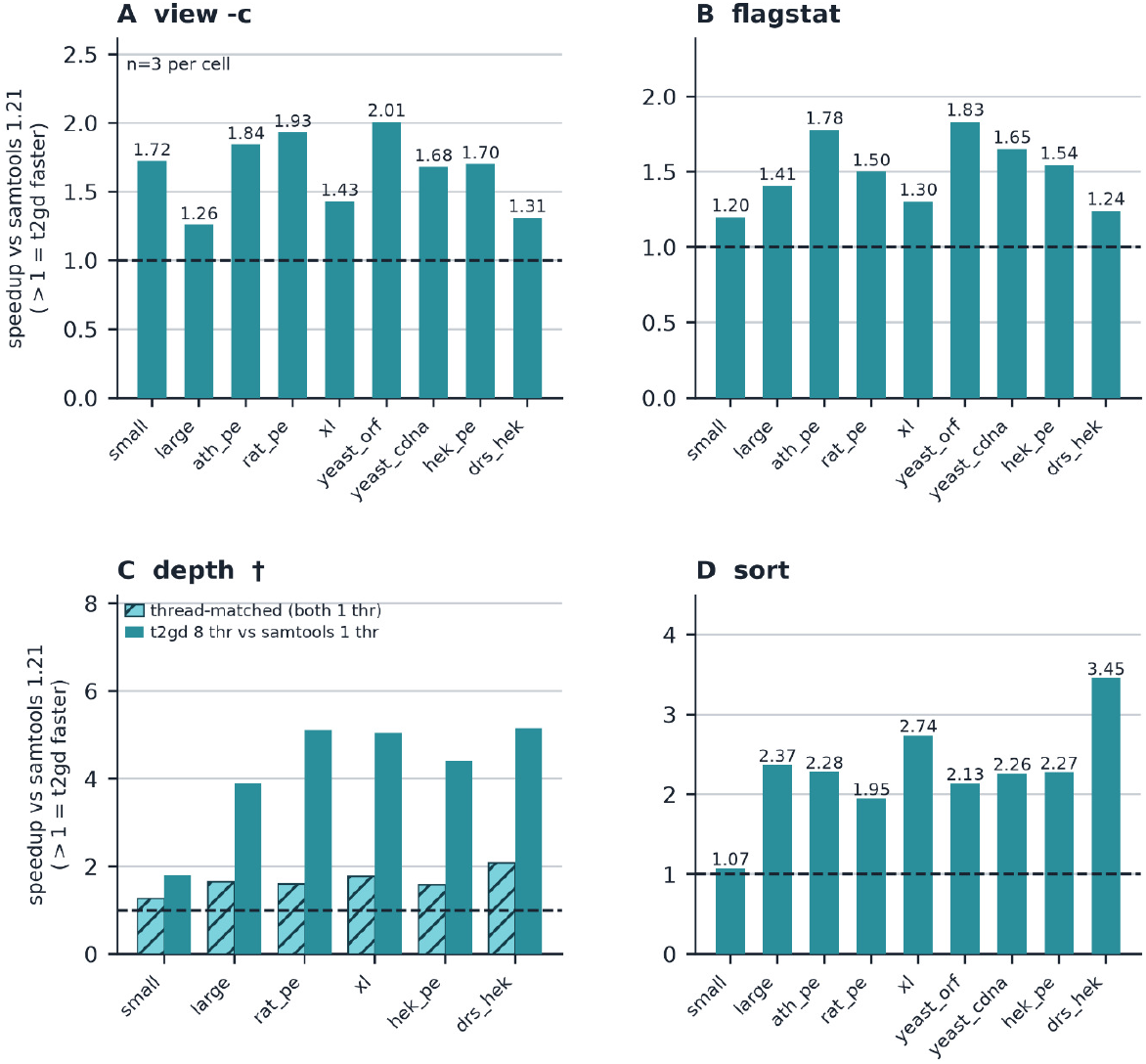
BAM-handling performance of t2gd against samtools. Speed-up of t2gd v1.19.1 over samtools 1.21 (*t*_samtools_*/t*_t2gd_; values above the dashed line at 1 favour t2gd) for (**A**) view -c, (**B**) flagstat, (**C**) depth and (**D**) sort, mean of 3 replicates per dataset. In A, B and D both tools used 8 threads. In C (^*†*^), samtools depth is single-threaded: solid bars compare t2gd at 8 threads with samtools, and hatched bars compare both tools at one thread.

## Discussion

Across long-read direct RNA sequencing, ultra-short ribosome footprinting and paired-end short-read RNA sequencing, aligning to the transcriptome and then lifting delivers its benefit through one mechanism: splice-junction and pseudogene placement is resolved deterministically from the annotation rather than inferred from the read alone. The same mechanism is the source of the limitations. Restricted to annotated transcript sequence, T2G cannot detect novel splice junctions, unannotated isoforms or intron-retention signal. T2G should therefore be run alongside a genome-direct pipeline, not instead of one, whenever novel splicing or transcription is itself of interest. Within that scope it returns transcriptome resolution to genomic coordinates without discarding the CIGAR structure or the tag payload that make the alignments worth keeping, and packages the result, together with quantification, metadata, and BAM handling utilities, as a single dependency-free binary.

## Data availability

The direct RNA sequencing data generated for this study are deposited in the NCBI Sequence Read Archive under BioProject accession PRJNA1512710. All other datasets are previously published and publicly available: paired-end RNA sequencing under SRA accessions SRR31348820 and SRR31348821; ribosome footprinting under SRR493747; the scalability panel under SRR31348821 (human), SRR33313115 (rat), SRR18592149 (yeast) and ENA accession ERR16142461 (*A. thaliana*); and POND-seq under SRR37350470. Reference sequences and annotations are from Ensembl release 115 (human GRCh38, rat GRCr8, *S. cerevisiae* R64-1-1), Ensembl Plants release 62 (*A. thaliana* TAIR10/Araport11) and the Saccharomyces Genome Database (*S. cerevisiae* S288C R64-5-1, release 20240529).

## Code availability

T2G is released under CC BY-NC-ND 4.0. Documentation, pre-built binaries for Linux x86 64 (broadwell, znver2, znver4), macOS arm64 and Windows x86 64 (portable archive and installer), and the Python reference implementation (under python/) are hosted at https://github.com/Arnaroo/T2GD. The v1.19.1 snapshot, comprising the documentation and the Python reference, is archived at https://doi.org/10.5281/zenodo.22016829; the concept DOI https://doi.org/10.5281/zenodo.21744718 resolves to the current version. The tool is also available at https://biocodecs.org/tools. The reference implementation requires Python 3 and pysam and is published so that the method is auditable; the optimised implementation is distributed as a compiled binary and its D source tree is not part of the release. Benchmark scripts, summary data and figure-rendering scripts are attached to the v1.19.1 release as T2GD supplementary code and data.tar.gz and described under reproducibility/ in the repository.

## Acknowledgments

The authors thank the Biocodecs Group, the RNA Innovation Foundry (RIF) and The Australian Centre for RNA Therapeutics in Cancer (ACRTC) teams for discussions throughout the project, and acknowledge use of the Kaya HPC facility at UWA together with the support of the Kaya HPC team, in particular Chris Bording.

